# Anatomy of the Visual Word Form Area in Dyslexia

**DOI:** 10.64898/2026.07.31.742142

**Authors:** Jamie L. Mitchell, Maya Yablonski, Mia Jimenez, Howard Chiu, Jason D. Yeatman

## Abstract

The Visual Word Form Area (VWFA), located in ventral occipitotemporal cortex, plays a critical role in skilled reading. Researchers have theorized that the VWFA develops in its specific anatomical location due to the convergence of major white matter tracts and proximity to functionally similar regions. This suggests that precise anatomical positioning may be crucial for optimal VWFA function. Previous research has identified several functional differences in this region between typical and struggling readers (i.e. dyslexia): struggling readers show weaker text-selective responses and often exhibit a smaller or even absent VWFA. However, it remains unexplored whether the precise anatomical location of this region also differs between typical and struggling readers. We tested whether VWFA anatomy differs between children with and without dyslexia (N=87). Participants completed a functional localizer, which we used to manually define the VWFA in each individual’s native anatomy. We examined whether: (1) VWFA anatomical location relates to reading ability, (2) children with dyslexia show greater variability in VWFA location compared to typical readers, and (3) VWFA location with respect to white matter tracts relates to reading ability. Results reveal that, despite being smaller in children with dyslexia, there is no relationship between VWFA location and reading ability. Specifically, individual VWFA location relative to anatomy, relative to other’s VWFAs, and relative to white matter tracts, is not related to reading ability. These findings suggest that while the VWFA’s general anatomy may be facilitated by development, its precise location remains stable and unrelated to reading proficiency.

## 1 Introduction

The Visual Word Form Area (VWFA) is a high-level visual region that is at the foundation of rapid and automatic word recognition (Cohen et al., 2000, 2002). VWFA develops as multiple text-selective patches (Woodhead et al., 2014; Yeatman et al., 2014; Yeatman & White, 2021) in high-level visual cortex, specifically on the fusiform gyrus and along the occipitotemporal sulcus within ventral occipitotemporal cortex (VOTC; (Cohen & Dehaene, 2004; Grill-Spector & Weiner, 2014; McCandliss et al., 2003). These patches are often separated into a more posterior VWFA-1 and a more anterior VWFA-2 (Lerma-Usabiaga et al., 2018; White et al., 2019) as they exhibit variability in structural and functional underlying properties (Lerma-Usabiaga et al., 2018; Weiner et al., 2014). The development of VWFA is thought to be due to the change in category-specific tuning (like increased sensitivity to text and decreased sensitivity objects) throughout high-level visual cortex that occurs with reading experience and through development (Dehaene & Cohen, 2007; Kubota et al., 2024; Nordt et al., 2021). However, despite the relative anatomical consistency at a coarse spatial scale, the exact location of this region is highly variable (Glezer & Riesenhuber, 2013; Mitchell et al., 2026).

Experiential factors play an important role in the specialization of this region. Researchers have determined that the emergence of VWFA co-occurs with the onset of literacy acquisition and that it plays a key role in reading development (Ben-Shachar et al., 2011; Dehaene et al., 2010; Dehaene & Cohen, 2011; Dehaene-Lambertz et al., 2018; Li et al., 2023). This relationship between VWFA development and reading experience is emphasized in research surrounding dyslexia in which differential or even decreased activation of this region is observed in children with dyslexia (Dębska et al., 2021; Eden et al., 2004; Maisog et al., 2008; Mitchell et al., 2025; Shaywitz et al., 2002; van der Mark et al., 2009).

Text-selective tuning of VOTC is also thought to be the consequence of this region’s distinct structural (Bouhali et al., 2014) and functional (Yablonski et al., 2024) connectivity to language processing regions. Many researchers suggest that VWFA emerges in this area of visual cortex due to its proximal location to major white matter tracts that connect the visual and language processing regions of the brain (Bouhali et al., 2014; Caffarra et al., 2021; Yeatman et al., 2013; Yeatman & White, 2021). In fact, bundles such as the Inferior Longitudinal Fasciculus (ILF), Vertical Occipital Fasciculus (VOF), Arcuate (Arc) and posterior Arcuate (pArc) fascicles, and the Inferior Fronto-Occipital Fasciculus (IFOF) have been identified as key structures of the reading circuitry which intersect with this region of VOTC (Grotheer et al., 2021; Yeatman, Dougherty, Ben-Shachar, et al., 2012; Yeatman et al., 2013).

Research on reading related white matter development indicates that these structural pathways may be utilized differently prior to literacy development (Vandecruys et al., 2024) and that white matter cortical endpoint connectivity to VOTC is more categorically-distinct in adults than in children (Kubota et al., 2023). Furthermore, changes in tissue property of these fascicles has been shown to be related to targeted intervention and improved reading ability (Huber et al., 2018; Meisler et al., 2024).

Despite all this knowledge of the role VWFA plays in reading and the relationships that exist between reading ability and functional properties of VWFA, little is known about the role that the anatomy of this region plays in reading. While it is clear that VWFAs are smaller and less selectively tuned to text in dyslexic compared to typical readers (Mitchell et al., 2025), it is unclear if these differences are solely due to functional properties of the region, or if the location in which the region develops also plays a role. Could these differences be the result of VWFA emerging in a “sub-optimal” location? Specifically, we address 3 hypotheses linking the anatomy of the VWFA to reading skills with a focus on understanding the root of reading struggles. We hypothesize that in children with dyslexia versus typical readers, VWFA:

1. Appears in different anatomical locations
2. Location is be more variable
3. White-matter connection patterns differ

To investigate these hypotheses we take several approaches to measuring the anatomical location of VWFA, utilizing functional MRI (fMRI), diffusion MRI (dMRI), and reading assessments in a cohort of children consisting of typical readers and children with dyslexia. With this multimodal approach, we are able to explore the anatomy of VWFA at an individual level leading to a more detailed understanding of the link between anatomy, function, and the process of learning to read.

## 2 Methods

All study protocols were approved by Stanford University’s School of Medicine Institutional Review Board (IRB). Researchers obtained both written and verbal assent from participants and consent from at least one of the child’s parents or guardians. Participants were compensated a flat rate of $120 for study visits with no loss of compensation for uncompleted tasks. Furthermore, participants were incentivized to maintain effort and focus during tasks with small prizes.

### 2.1 Participants

The study included 87 enrolled participants, however only 84 participants (ages 7-13) were included in analyses as 3 participants had unusable functional data due to high motion. Participants were divided into two groups: struggling readers (referred to as dyslexia readers; N=63) and typical readers (N=24). Group designations were determined through a screening process which occurred prior to the start of the study. For more information on screening and group designations see Mitchell et al., 2025. All participants reported normal or corrected-to-normal vision and had no reported neurological or hearing impairments. Furthermore, all participants reported having learned English before the age of three and were either monolingual English speakers or reported using English for at least 60% of their daily communication. For more demographic information, see Supplementary Table S1.

### 2.2 Reading assessments

Reading ability was determined using the Woodcock-Johnson Basic Reading Skills (WJ BRS; (Schrank & Wendling, 2018) composite score. This assessment consists of two subtests - Word Attack and Letter Word Identification - which focus on measuring a student’s decoding and word reading skills. For more information on assessment administration, see Mitchell at al., 2025.

### 2.3 MRI Acquisition

Functional MRI data for this study were reported in Mitchell et al., 2025. Briefly summarizing, functional data was collected using a gradient EPI multiband sequence with whole brain coverage. The acquisition parameters included a TR of 1.19 s, a TE of 30 ms, and a flip angle of 62, resulting in a spatial resolution of 2.4 mm³ isotropic voxels. Please see the original manuscript for full details.

Diffusion MRI data for this study was collected with 180 diffusion-weighted volumes, distributed across three shells with the following parameters: 30 directions with b = 1000 s/mm2, 60 directions with b = 2000 s/mm2, and 90 directions with b = 3000 s/mm2, as well as 14 reference volumes without diffusion weighting (b = 0 s/mm2). A TR of 3335 ms with an echo time of 87 ms was used to collect 1.5mm^3^ isotropic voxels in 84 axial slices. This was achieved using a hyperband acceleration with a slice acceleration factor of 4. The scan duration totaled 11 minutes and was broken down to two 5.5-minute scans to allow children a short break. An additional scan of six non-diffusion-weighted volumes with a reversed phase encoding direction and the same parameters was also acquired to correct for echo-planar imaging (EPI) distortions during processing see Yablonski et al., 2025 for more details.

### 2.4 MRI Data Preprocessing

Diffusion data were pre-processed using the default pipeline in QSIprep 0.22.0 (Cieslak et al., 2021). Reconstruction was performed using QSIprep 0.22.0, which is based on Nipype 1.8.6 (K. Gorgolewski et al., 2011; K. J. Gorgolewski et al., 2018). Multi-tissue fiber response functions were estimated using the dhollander algorithm. FODs were estimated via constrained spherical deconvolution (CSD; (Tournier et al., 2004, 2008) using an unsupervised multi-tissue method (Dhollander et al., 2016, 2019). Reconstruction was done using MRtrix3 (Tournier et al., 2019). FODs were intensity-normalized using mtnormalize (Raffelt et al., 2017).Many internal operations of qsiprep use Nilearn 0.10.1 (Abraham et al. 2014, RRID:SCR_001362) and Dipy 1.8.0(Garyfallidis et al. 2014).

Functional MRI data and T1w images were preprocessed using fMRIprep as in Mitchell et al., 2025. Of note, T1w images were first preprocessed with QSIprep before fmriprep and freesurfer preprocessing occurred. As such, care was taken to ensure that functional and diffusion data were properly aligned and registered to the same space.

#### 2.4.1 Fiber Tractography

pyAFQ 3.3 (Kruper et al., 2021, 2025; Yeatman, Dougherty, Myall, et al., 2012) was used on the whole-brain tractograms from the QSIprep pipeline to segment white matter tracts of interest and then calculate streamline density maps for each segmented white matter tract. Streamline density maps were then projected to native surface for each individual.

### 2.5 Region of Interest Definition

Functional regions of interest (ROI), as reported in Mitchell et al., 2025, were defined on the native surface for each individual. In the present study, all analyses regarding ROI location were performed in the fsaverage template space. To accomplish this, natively-defined ROIs were projected to the average surface. Furthermore, ROI centers were calculated using these fsaverage labels.

Individual-level ROI centers were calculated by finding the single vertex contained within the label that minimized the euclidean distance to all other vertices contained within the label. Group-level ROI centers were calculated in a similar manner in that a single participants center vertex was selected that minimized the euclidean distance to all other participants center vertex.

### 2.6 Statistical Analyses

All statistical analyses (unless otherwise specified) were performed in Python using scipy.stats and numpy libraries (Harris et al., 2020; Virtanen et al., 2020). Two-sided statistical tests were used throughout, with a significance threshold of *α* = 0.05.

Group differences in VWFA size, location, and variability were assessed using Welch’s t-tests, to account for unequal variances between groups. For analyses involving multiple comparisons (VWFA-1 and VWFA-2 across x, y, and z coordinates), Bonferroni correction was applied to adjust thresholds (adjusted *α* = 0.05/6 = 0.0083).

Relationships between continuous variables VWFA measures (i.e., size, location, variability) and reading ability (WJ BRS scores) were evaluated using Pearson correlations.

For all instances of null findings, Bayes factor was calculated to quantify the support for the null hypothesis over the alternative hypothesis. Bayes factors are reported as *bf_01_*, indicating the ratio of the likelihood that the data can be observed under the null hypothesis. Bayes factors were calculated with the Pingouin (Vallat, 2018) python package. For *bf_01_* calculations on t-tests, a Cauchy prior of 0.707 was used, and for Pearson correlations, the default uniformly stretched-Beta prior was used.

## 3 Results

### 3.1 VWFA Size is Related to Reading Ability

In our previous work (see Mitchell et al., 2025) we found that the size of Visual Word Form Area (VWFA) is smaller in children with dyslexia compared to typical readers (Figure1a-b). Furthermore, we demonstrated that VWFA is related to a child’s reading ability regardless of dyslexia status by comparing their VWFA size to their Woodcock-Johnson Basic Reading Skills scores (WJ BRS; Figure 1c). In order to confirm this size difference we again calculated the size of VWFA in terms of total cortical surface area devoted to each region of interest. In doing so, we find that VWFA-1 is, on average, 27.213 *mm^2^*(*sd* = 48.261) in participants with dyslexia and 89.580 *mm^2^*(*sd* = 95.042) in typically reading participants. We also found that VWFA-2 is, on average, 29.312 *mm^2^*(*sd* = 42.035) in dyslexic participants and 148.640 *mm^2^*(*sd* = 152.810) in typically reading participants. This indicates that in dyslexic participants, the total cortical surface area devoted to text-recognition in VWFA-1 is roughly 30% the size of a typical reader (*d* = -0.946) and WVFA-2 is roughly 20% the size of a typical reader (*d* = -1.324).

**Figure 1.**
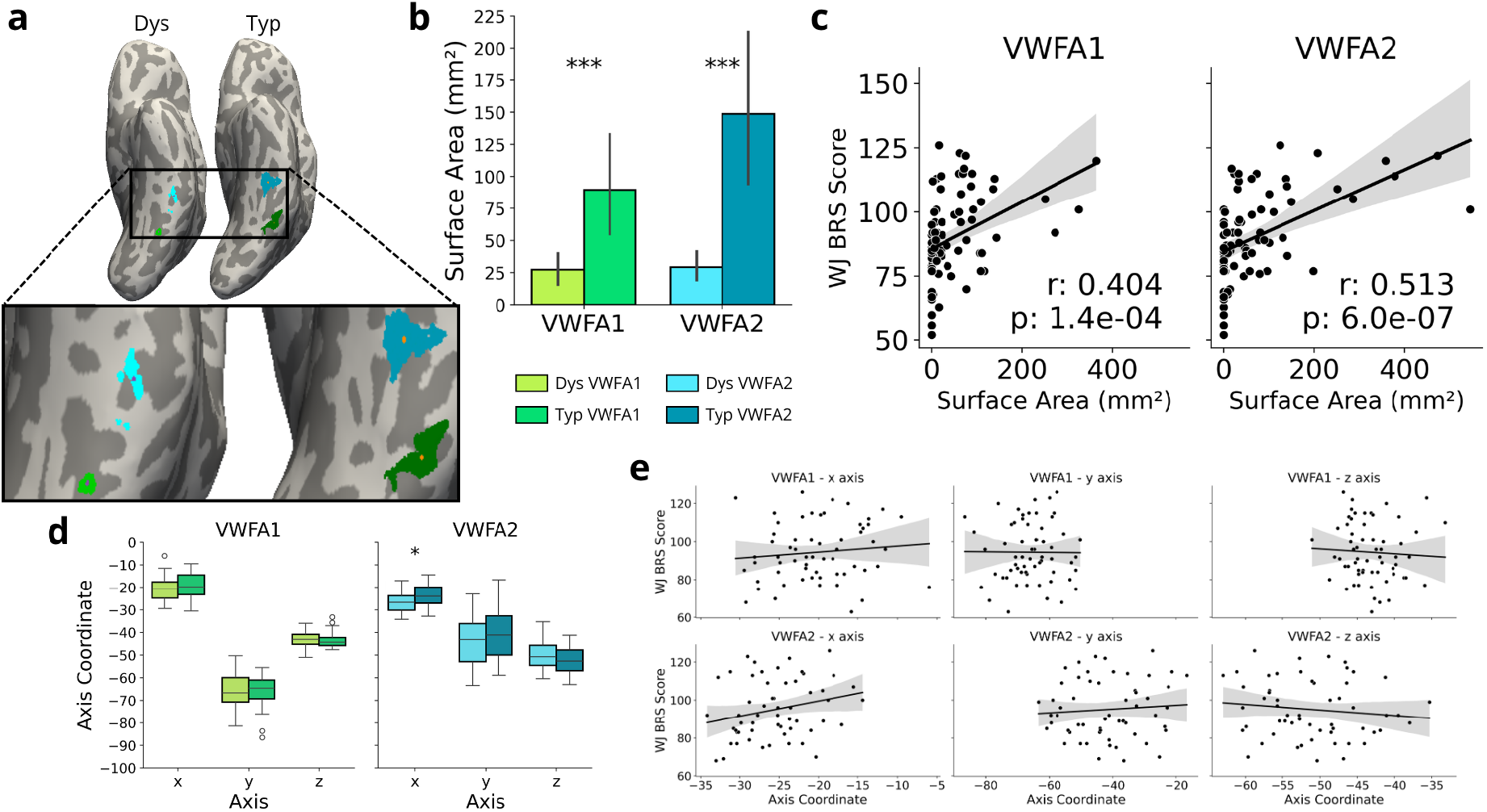
Location of the VWFA. **a** Sample VWFA-1 (greens) and VWFA-2 (blues) projected on the inflated cortical surface of one typical (darker shades) and one dyslexic participant (lighter shades). Orange dots on the typical participant and purple dots on the dyslexic participant represent the center of each VWFA. **b** VWFA size (in surface area, mm^2^) in typical readers (n = 24) and children with dyslexia (n = 60) for each region of interest (ROI). **c** Relationship between VWFA size and Woodcock-Johnson Basic Reading Skill score (WJ BRS). **d** Locations of center points of VWFAs 1 & 2 separated by reading group for x, y, and z MNI coordinates. Box plots display the median indicator, the box displays the interquartile range (IQR; 25%-75% range), and whiskers represent +/− 1.5 times the IQR. Outliers in the data are individually plotted above and below box whiskers. **e** Relationship between VWFA center point MNI Coordinates and WJ BRS. All results are unrelated (all results p > 0.05). Asterisks in **b** and **d** indicate the degree of significance derived from a two-sides Welch’s t-test (p < 0.001: ***, p < 0.01: **, p < 0.05: * ) Error bars in **b** and error bands in **c** and **e** represent a 95% confidence interval. Data for **b** and **c** are taken from Mitchell et al., 2025.

### 3.2 VWFA Location is Unrelated to Reading Ability

Motivated by the findings described above, we next sought to investigate our first hypothesis to determine if the location of VWFA was related to reading ability. The stark difference in size of VWFA in both groups led us to believe that there may be other physical differences of VWFA. The size of VWFA reflects the extent of cortical territory that has become selectively responsive to text, while the location reflects where that specialization emerges. That is, two individuals may have VWFAs of comparable sizes, while those regions occupy very different portions of the ventral occipitotemporal cortex. The precise location at which text selectivity emerges in an individual may therefore relate to behavioral characteristics.

To investigate this, we first found the center of each VWFA (see Methods) and recorded the corresponding MNI coordinates for each participant. We then compared the two groups of participants by running a Welch’s *t* test comparing each coordinate along the x, y, and z axes for both VWFA-1 and VWFA-2 (Figure 1d). We found that there were no group differences in x (*t*(44.98) = -0.920; *p* = 0.3625; *CI* = (-4.21, 1.57); *bf_01_*= 2.627), y (*t*(49.40) = 0.435; *p* = 0.6654; *CI* = (-3.33, 5.17); *bf_01_* = 3.455), or z (*t*(41.44)=0.169 (p=0.8665; CI=(-1.75, 2.07); *bf_01_* = 3.695) MNI coordinates for VWFA-1. We also found no group differences in the y (*t*(44.34) = -0.947; *p* = 0.3488; *CI* = (-9.71, 3.50); *bf_01_* = 2.509) and z (*t*(49.20) = 1.551; *p* = 0.1273; *CI* = (-0.78, 6.02); *bf_01_* = 1.355) MNI coordinates for VWFA-2. We did find a slight difference in the x-axis location of VWFA-2 between the groups (*t*(39.20) = -2.308; *p* = 0.0264; *CI* = (-5.48, -0.36); *bf_01_* = 0.422) suggesting that dyslexic participants have a slightly more lateral VWFA-2 compared to typical participants. However, this result did not survive a Bonferroni correction accounting for multiple comparisons (2 ROIs * 3 axes) (p = 0.158).

To go beyond a binary distinction, we tested the relationship between each participant’s coordinate locations and their reading score as a continuous variable, for both VWFAs (Figure 1e). A Pearson correlation revealed that there is indeed no relationship between reading score and the position of VWFA-1 along all axes nor the position of VWFA-2 along the z and y axes (all *p* > 0.3, *bf_01_* > 4, see Figure 1e), as seen previously with *t*-test results. Furthermore, the previously observed borderline relationship between the x coordinate location and dyslexia in VWFA-2 was statistically insignificant (*p* = 0.071, *bf_01_* = 1.223) when examined on this continuous scale correlated to reading score. Combined, these results show that the location of the center of VWFAs 1 & 2 is unrelated to a child’s reading ability.

### 3.3 Between-Participant Variability in VWFA Location is Unrelated to Reading Ability

Despite the lack of evidence for location-specific relationships to reading ability, the extent of variability observed amongst precise VWFA locations was vast enough that we suspected the variability may play a role in behavioral outcomes of reading. Researchers suggest the VWFa emerges in an optimal anatomical location for visual word processing (Hasson et al., 2002; Malach et al., 2002), so we suspect that certain, fine-grained locations may specialize for text-processing more frequently in typical readers compared to those with dyslexia.

Specifically, we hypothesized that children with dyslexia would have greater variability in VWFA location (i.e., less consistent location) compared to typical readers. Furthermore, quantifying variability is particularly important in dyslexia because greater spatial heterogeneity could help explain why group averaged studies sometimes report weak or inconsistent VWFA responses (Glezer & Riesenhuber, 2013; Mitchell et al., 2026).To test this, we first determined a single point of comparison for each group and VWFA label. Using the center points from each individual’s VWFA, we found a group center by selecting the center point from one individual whose surface vertex minimized the total distance to all other individuals center points for each group of participants (Figure2a). In doing so, we created a group center for VWFAs 1 & 2 for both groups.

Upon visual inspection, we observed little difference in the spread of the group centers (Figure 2b). To quantify this relationship, we used the group centers for each VWFA to calculate the average distance from each individual’s VWFA center to the group center. Using these distances as an indicator of the average spread of VWFAs, we then directly compared these distances between groups with a Welch’s *t*-test (Figure 2c). In doing so, we confirmed there was no difference between dyslexic and typical readers in the spread of centers in both VWFA-1 (*t*(42.85)=0.245; p=0.808; *CI*=(-2.73, 3.49); *bf_01_* = 3.647) and VWFA-2 (*t*(45.99)=-0.109; *p*=0.913; *CI*=(-4.04, 3.62); *bf_01_* = 3.616).

**Figure 2.**
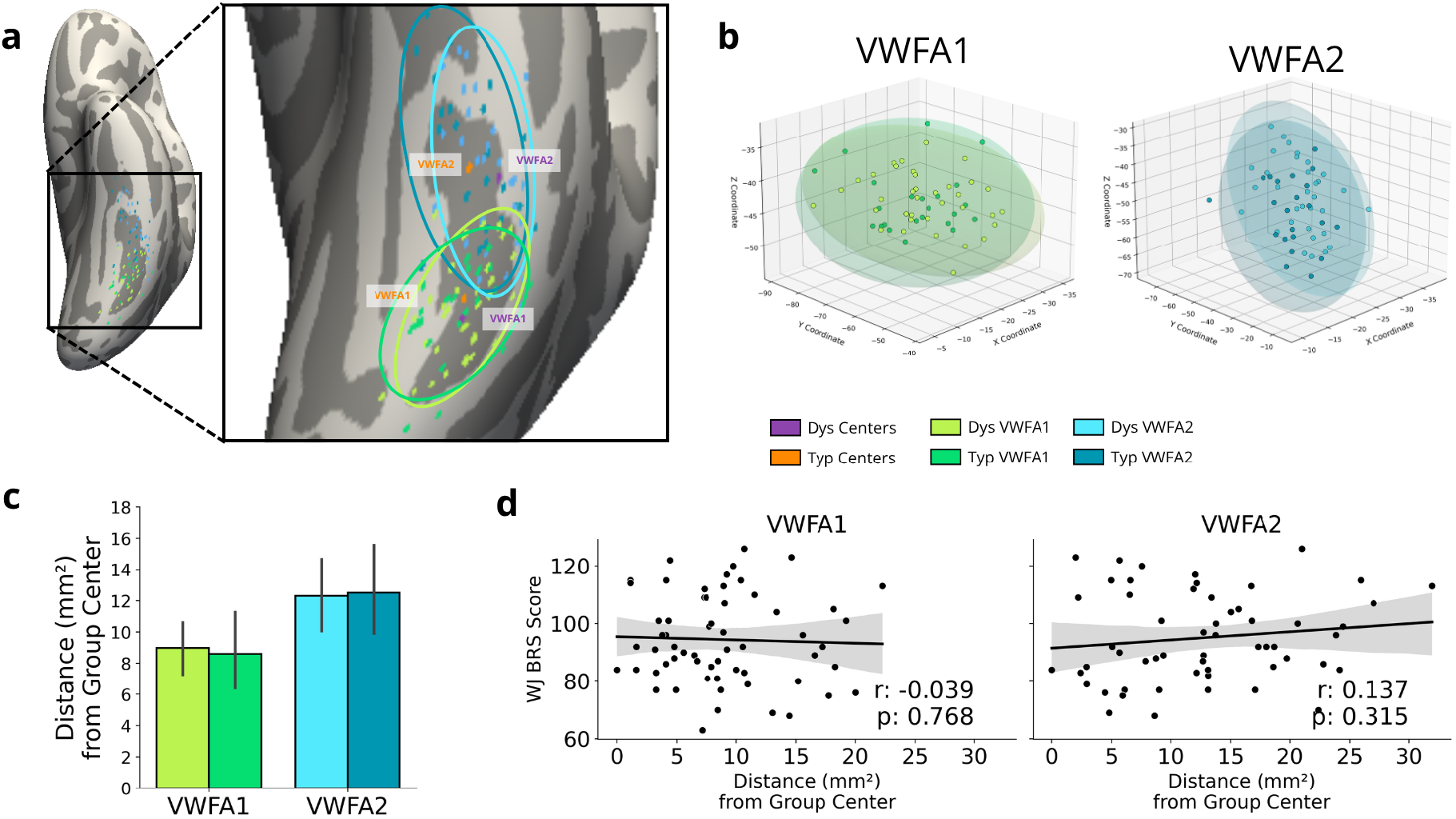
Variability in VWFA Location. **a** VWFA-1 (greens) and VWFA-2 (blues) centers in typical (darker shades) and dyslexic (lighter shades) participants projected on an inflated cortical surface in template space. Centers are defined as the single vertex that minimizes the distance between all vertices in the VWFA label. Orange dots represent the group-level centers of both VWFAs in typical participants and purple dots represent the group-level centers of both VWFAs in dyslexic participants. Ellipses display the approximate 95% confidence interval of the x and y MNI coordinates for each VWFA and group. **b** 3-dimensional MNI coordinates for each VWFA and group. Ellipsoids display the actual 95% confidence interval region for each VWFA and group. **c** Average spread of VWFAs in typical readers and children with dyslexia. Spread is calculated as the Euclidean distance between each individual’s VWFA center and the group center. **d** Relationship between individual VWFA distance to the all-subject VWFA center and WJ BRS. Error bars in **c** and error bands in **d** represent a 95% confidence interval.

We again sought to determine if there is a relationship between distance and reading ability on a continuous scale. To accomplish this, we first calculated a center point for both VWFA-1 and VWFA-2, this time using the centers across all participants’ respective VWFA. We then ran a Pearson correlation comparing each participants’ WJ BRS score to the distance between each participant’s ROI center and this new group-level center (Figure 2d). We found no significant relationship for both VWFA-1 (*r* = -0.039, *p* = 0.768, *bf_01_* = 6.00) and VWFA-2 (*r* = 0.137, *p* = 0.315, *bf_01_* = 3.67). Combined with the previously reported binary group-level results, these findings suggest that individual-level variability in VWFA location is not significantly related to reading ability.

### 3.4 Structural Connectivity of VWFA is Unrelated to Reading Ability

Size, location, and location variability characterize VWFA as a cortical region, but reading isn’t supported by VWFA in isolation. Fluent reading requires visual information to integrate with distributed systems that support various other processes (Posner & McCandliss, 1999; Price, 2000). Structural connectivity therefore provides a complementary investigative approach to understanding the underlying anatomy of VWFA - whereas the previous analyses characterize VWFA as a specific region, structural connectivity characterizes the existing connections to the region that facilitate the process of reading. Therefore, we suspect that the underlying connections of VWFA may differ in children with dyslexia compared to typical readers despite lack of evidence in differences of cortical location. To test our third hypothesis, we used diffusion MRI to examine white matter tracts connected to the VWFA and determine whether structural connectivity patterns of the VWFA relate to reading ability.

For this investigation, we included streamline data from six major white matter tracts (Grotheer et al., 2021; Yeatman, Dougherty, Ben-Shachar, et al., 2012; Yeatman et al., 2013) that are involved with reading (Figure 3a); the Arcuate Fasciculus (Arc), Posterior Arcuate Fasciculus (pArc), Inferior Fronto-Occipital Fasciculus (IFOF), Inferior Longitudinal Fasciculus (ILF), Anterior Vertical Occipital Fasciculus (aVOF), and Posterior Vertical Occipital Fasciculus (pVOF). Each participant’s streamline density maps generated from the pyAFQ (see methods; (Garyfallidis et al., 2014, 2018; Kruper et al., 2021, 2025; Yeatman, Dougherty, Myall, et al., 2012) were first projected to the cortical surface (Figure 3b). We then extracted the average bundle density (reported as number of streamlines per vertex) within the VWFA of each participant. This gave us an estimate for the density of different white matter tracts that intersect with each participant’s VWFA (Figure 3c; (Kubota et al., 2023; Takemura et al., 2016). We first observed the expected pattern of distinct connectivity profiles for VWFA-1 and VWFA-2. Specifically, VWFA-1 intersects with the ILF and VOF, while VWFA-2 shows greater connectivity with the Arc, in line with previous findings (Grotheer et al., 2021; Kubota et al., 2023; Lerma-Usabiaga et al., 2018). We then ran a Welch’s *t*-test comparing the average fiber density within each VWFA and for each of the seven white matter bundles and found no significant difference in mean density for any of the fiber bundles in either of the VWFAs between dyslexic and typical readers (*p* > 0.1, *bf_01_* > 1.5; Supplementary Table S2).

**Figure 3.**
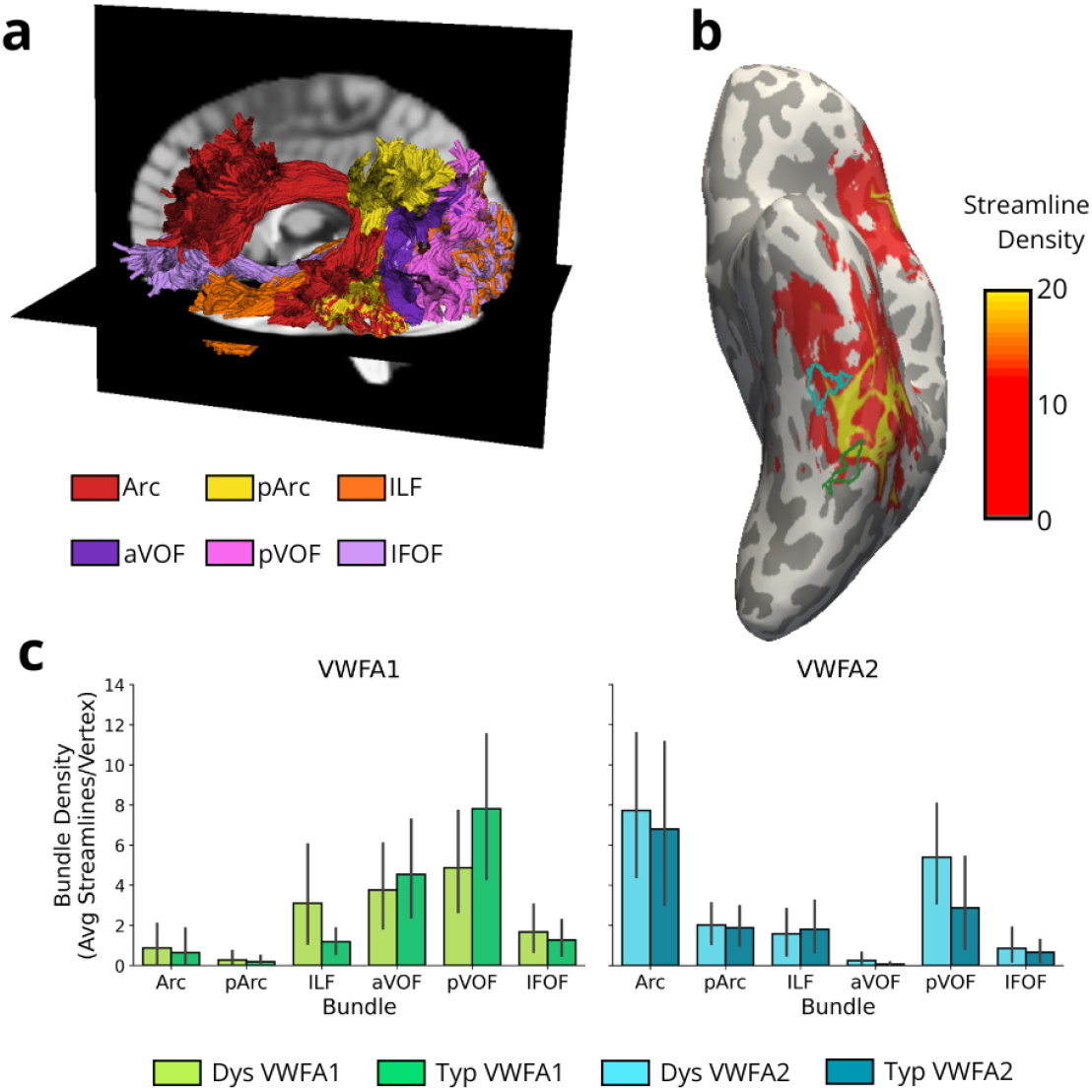
VWFA-Constrained White Matter Bundle Density. **a** White matter bundle streamlines displayed in the left hemisphere of the native T1w space for one example participant. Arcuate Fasciculus (red): Arc; Posterior Arcuate Fasciculus (yellow): pArc; Inferior Longitudinal Fasciculus (orange): ILF; Anterior Vertical Occipital Fasciculus (purple): aVOF; Posterior Vertical Occipital Fasciculus (pink): pVOF; Inferior Fronto-Occipital Fasciculus (light purple): IFOF. **b** Density map expressing the mean number of streamlines for the Arcuate Fasciculus for one representative participant, projected onto their fsnative left inflated cortical surface. VWFA-1 (green) and VWFA-2 (blue) contours are layered over the density map and demonstrate the process used to calculate average streamline density. **c** Average streamline density in the VWFA, for each white matter tract in participants with dyslexia (lighter shades) and in typical readers (darker shades). Error bars represent the standard error.

We then evaluated this relationship on a continuous scale using a Pearson correlation of each participant’s average streamline density and their WJ BRS score (Supplementary Table S3). This analysis also revealed no significant relationship between streamline density and WJ BRS performance for any white matter bundle in either VWFA-1 or VWFA-2 (p > 0.3) These results indicate that the VWFAs’ location relative to reading-related white matter tracts is generally significantly related to reading ability.

## 4 Discussion

Previous research has revealed that functional properties of the VWFA are significantly related to reading ability (Ben-Shachar et al., 2011; Dehaene et al., 2010; Dehaene & Cohen, 2011; Dehaene-Lambertz et al., 2018; Li et al., 2023). Dyslexia research has further emphasized this relationship (Dębska et al., 2021; Eden et al., 2004; Maisog et al., 2008; Shaywitz et al., 2002; van der Mark et al., 2009) and has revealed that structural changes can be observed in concert with improvements in reading ability following intensive intervention (Huber et al., 2018; Meisler et al., 2024). Our own previous work has shown that improvement in reading ability corresponds with an increase in size and selectivity of VWFA (Mitchell et al., 2025). Because previous work suggests that VWFA emerges in a location that is ideal for the processing of text (Hasson et al., 2002; Malach et al., 2002; Yeatman & White, 2021), we hypothesized that a suboptimal location could be related to the struggles children with dyslexia face with gaining proficiency. Consequently, this study sought to examine whether the anatomical location of the VWFA is related to reading ability. We found, however, that the location of VWFA relative to cortical folds and white matter tracts was not significantly related to reading ability.

A fundamental question in reading neuroscience is why the VWFA consistently emerges in ventral occipitotemporal cortex. One influential hypothesis suggests that this location is determined by its optimal connectivity profile—positioned at the intersection of visual processing streams and language networks, allowing efficient integration of orthographic information with phonological and semantic systems (Grotheer et al., 2021; Lerma-Usabiaga et al., 2018; Yeatman & White, 2021). If connectivity determines the optimal location for VWFA, then suboptimal positioning in children with dyslexia could lead to inefficient information transfer, potentially contributing to reading difficulties.

We used multimodal mri to look at structural connectivity of the functionally defined VWFA in individual children with a wide range of reading ability. .Our investigation started with the relationship between the position of the VWFA in relation to reading ability. We found that VWFA location on the cortical surface was not related to reading ability, contrary to our hypothesis. We then went on to determine if there was more variability in the location of VWFA for struggling readers with an assumption that perhaps typical readers may develop VWFA in a more consistent location than those with dyslexia. We again found no relationship. Combined, these two location-based findings suggest that exact cortical position of the VWFA may not be a determining factor in someone’s reading performance.

Examining structural connectivity moves the analysis from identifying where readers differ to investigating a possible mechanism through which these differences, if any, arise or affect reading. Previous research suggests that white matter tissue properties change with reading improvement (Huber et al., 2018; Meisler et al., 2024) and differential structural profiles of reading-related regions between children (i.e. less experienced readers) and adults (i.e. more experienced readers; (Kubota et al., 2023). Despite these these reading-related white-matter properties, we found no evidence of a direct relationship between streamline densities for major reading tracts that intersect with VWFA and one’s reading ability. This suggests that VWFA location relative to white matter tracts is not indicative of reading ability.

The lack of evidence of anatomical differences in VWFA for children with dyslexia indicates that the exact anatomy of the region, though highly variable between individuals, is not a biomarker of dyslexia. Instead, these results emphasize within the VWFA, functional properties , like tuning and the extent of text-selective activation of the region, are the primary biological correlates of reading ability. However, further research utilizing intervention approaches may reveal within-subject tendencies for anatomical changes of this region which may correspond to behavioral changes in reading performance.

## 5 Data and Code Availability

De-identified data has been made publicly available through the Stanford University Libraries Digital Repository and can be found at https://purl.stanford.edu/qq284rg5214

Code has been made publicly available through an online GitHub repository and can be found at https://github.com/jamielmitchell/Mitchell_VWFA-anatomy

## 6 Author Contributions

**JLM:** Conceptualization, Formal analysis, Investigation, Writing (original draft & review and editing), Visualization. **MY:** Investigation, Writing (review and editing). **MJ:** Investigation, Data curation, Project administration, Writing (review and editing). **HC:** Writing (review and editing). **JDY:** Conceptualization, Resources, Writing (review and editing), Supervision, Funding acquisition.

## Acknowledgements

We would like to thank the individuals and families who participated in this study. We would also like to thank the research coordinators and assistants who contributed to data collection and in particular would like to thank Megumi E. Takada for her contributions coordinating the larger study efforts.

