## Supplemental Material for "Anatomy of the Visual Word Form Area in Dyslexia"

### Anatomy of the Visual Word Form Area in Dyslexia: Supplemental Material

| Participant Information | | |
|---|---|---|
| | ***Dyslexic*** | ***Typical*** |
| N | 60 | 24 |
| Gender (f/m) | 28 / 31* | 13 / 11 |
| Mean age ± SD | 10.0 ± 1.3 | 9.9 ± 1.4 |
| Multilingual Count | 21 | 11 |
| WJ BRS | 81.5 ± 10.1 | 110.1 ± 8.3 |

**Table S1 | Participant Information**
Demographic information, Woodcock-Johnson Basic Reading Skills (WJ BRS) scores, and functional MRI task accuracy for participants divided by study group. Gender and multilingual status are collected as self-report responses. Age and task performance are displayed as the average ± standard deviation. * One participant self-reported gender as non-binary.

| White Mater Bundle Densities in VWFA | | | | | | | | |
|---|---|---|---|---|---|---|---|---|
| | **Dyslexia** | **Typical** | **t** | **CI** | | **DOF** | ***bf***$_{01}$ | ***p*** |
| | *Mean ± sd* | *Mean ± sd* | *(Dys > Typ)* | *low* | *high* | | | |
| **VWFA1** | | | | | | | | |
| Arc | 0.878 ± 3.106 | 0.635 ± 2.450 | 0.332 | -1.222 | 1.706 | 54.531 | 3.572 | 0.742 |
| pArc | 0.271 ± 1.294 | 0.179 ± 0.681 | 0.358 | -0.423 | 0.608 | 58.242 | 3.545 | 0.722 |
| IFOF | 1.673 ± 3.892 | 1.259 ± 2.193 | 0.523 | -1.171 | 2.000 | 58.846 | 3.336 | 0.603 |
| ILF | 3.101 ± 7.832 | 1.171 ± 1.577 | 1.451 | -0.755 | 4.616 | 41.893 | 1.562 | 0.154 |
| aVOF | 4.869 ± 7.694 | 7.804 ± 8.603 | -1.317 | -7.431 | 1.561 | 42.222 | 1.818 | 0.195 |
| pVOF | 3.766 ± 6.810 | 4.531 ± 6.154 | -0.443 | -4.229 | 2.700 | 49.955 | 3.444 | 0.659 |
| **VWFA2** | | | | | | | | |
| Arc | 7.717 ± 10.692 | 6.796 ± 9.453 | 0.331 | -4.664 | 6.505 | 48.611 | 3.472 | 0.742 |
| pArc | 2.016 ± 2.920 | 1.877 ± 2.380 | 0.191 | -1.320 | 1.598 | 50.824 | 3.579 | 0.849 |
| IFOF | 0.855 ± 2.533 | 0.665 ± 1.299 | 0.363 | -0.862 | 1.242 | 51.958 | 3.441 | 0.718 |
| ILF | 1.582 ± 3.559 | 1.803 ± 3.077 | -0.241 | -2.056 | 1.616 | 49.257 | 3.547 | 0.811 |
| aVOF | 5.401 ± 7.359 | 2.867 ± 5.498 | 1.444 | -0.987 | 6.055 | 52.639 | 1.543 | 0.155 |
| pVOF | 0.250 ± 0.999 | 0.067 ± 0.208 | 1.018 | -0.181 | 0.547 | 37.365 | 2.369 | 0.315 |

**Table S2 | White Matter Bundle Densities**

Results from a Welch's *t*-test between the bundle densities of seven reading-related white matter tracts in VWFAs of dyslexic and typical readers. Bundle density is calculated as the mean number of streamlines per vertex. Arcuate Fasciculus (Arc), Posterior Arcuate Fasciculus (pArc), Temporoparietal Fasciculus (TP), Inferior Fronto-Occipital Fasciculus (IFOF), Inferior Longitudinal Fasciculus (ILF), Anterior Vertical Occipital Fasciculus (aVOF), Posterior Vertical Occipital Fasciculus (pVOF). Significant results are displayed in bold and asterisks indicate the degree of significance ($p < 0.001$: ***, $p < 0.01$: **, $p < 0.05$: *)

| Bundle Density & Reading Score Correlations | | | | | | |
|---|---|---|---|---|---|---|
| | r | CI | | DOF | $bf_{01}$ | p |
| | | *low* | *high* | | | |
| **VWFA1** | | | | | | |
| Arc | -0.055 | -0.303 | 0.199 | 59 | 5.735 | 0.673 |
| pArc | 0.001 | -0.251 | 0.253 | 59 | 6.257 | 0.994 |
| IFOF | -0.062 | -0.309 | 0.193 | 59 | 5.605 | 0.635 |
| ILF | -0.117 | -0.359 | 0.139 | 59 | 4.211 | 0.368 |
| aVOF | 0.128 | -0.127 | 0.368 | 59 | 3.889 | 0.324 |
| pVOF | 0.024 | -0.229 | 0.274 | 59 | 6.154 | 0.854 |
| **VWFA2** | | | | | | |
| Arc | 0.016 | -0.248 | 0.277 | 54 | 5.959 | 0.909 |
| pArc | 0.024 | -0.241 | 0.285 | 54 | 5.912 | 0.863 |
| IFOF | 0.086 | -0.181 | 0.341 | 54 | 4.938 | 0.528 |
| ILF | 0.134 | -0.134 | 0.383 | 54 | 3.742 | 0.325 |
| aVOF | -0.081 | -0.336 | 0.186 | 54 | 5.057 | 0.554 |
| pVOF | 0.006 | -0.257 | 0.269 | 54 | 5.991 | 0.962 |

**Table S3 | White Matter Bundle Density and Reading Ability**

Results from a Pearson Correlation between white matter bundle density of a participant's VWFA and their WJ BRS (Woodcock-Johnson Basic Reading Skills) score. Bundle density is calculated as the mean number of streamlines per vertex. Arcuate Fasciculus (Arc), Posterior Arcuate Fasciculus (pArc), Temporoparietal Fasciculus (TP), Inferior Fronto-Occipital Fasciculus (IFOF), Inferior Longitudinal Fasciculus (ILF), Anterior Vertical Occipital Fasciculus (aVOF), Posterior Vertical Occipital Fasciculus (pVOF). Significant results are displayed in bold and asterisks indicate the degree of significance ($p < 0.001$: ***, $p < 0.01$: **, $p < 0.05$: *)
